# A rigorous measure of genome-wide genetic shuffling that takes into account crossover positions and Mendel’s second law

**DOI:** 10.1101/194837

**Authors:** Carl Veller, Nancy Kleckner, Martin A. Nowak

## Abstract

Comparative studies in evolutionary genetics rely critically on evaluation of the total amount of genetic shuffling that occurs during gamete production. However, such studies have been hampered by the fact that there has been no direct measure of this quantity. Existing measures consider crossing over by simply counting the average number of crossovers per meiosis. This is qualitatively inadequate because the positions of crossovers along a chromosome are also critical: a crossover towards the middle of a chromosome causes more shuffling than a crossover towards the tip. Moreover, traditional measures fail to consider shuffling from independent assortment of homologous chromosomes (Mendel’s second law). Here, we present a rigorous measure of genome-wide shuffling that does not suffer from these limitations. We define the parameter *r̅* as the probability that the alleles at two randomly chosen loci will be shuffled in the production of a gamete. This measure can be decomposed into separate contributions from crossover number and position and from independent assortment. Intrinsic implications of this metric include the fact that *r̅* is larger when crossovers are more evenly spaced, which suggests a novel selective advantage of crossover interference. Utilization of *r̅* is enabled by powerful emergent methods for determining crossover positions, either cytologically or by DNA sequencing. Application of our analysis to such data from human male and female reveals that: (i) *r̅* in humans is close to its maximum possible value of 1/2, (ii) this high level of shuffling is due almost entirely to independent assortment, whose contribution is ~30 times greater than that of crossovers.

## Introduction

The shuffling of maternally and paternally inherited genes during gamete production is an important process in sexual populations [1]. It improves the efficiency of adaptation by allowing natural selection to act separately on distinct mutations [2–7], and has been implicated in protecting populations from rapidly evolving parasites [8, 9] and from the harmful invasion of selfish genetic complexes [10–13]. The total amount of shuffling that occurs in gamete production is therefore a quantity of considerable importance, and has been the subject of much empirical interest.

Correspondingly, comparative studies of genome-wide shuffling have been carried out across species ([1, 14–20]; reviewed in [21–23]), with implications ranging from distinguishing the evolutionary advantages of sex [1, 14, 22–24] to testing the genomic effects of domestication [14, 16]. A large literature has also studied male/female differences in shuffling [13, 25, 26], prompting several evolutionary theories to explain these differences [12, 13, 25–28], which could concomitantly be informative of the adaptive value of shuffling [25]. There can also exist differences in shuffling across individuals of the same sex (e.g., in flies [29–33], mice [34–36], humans [36–41], and *Arabidopsis* [42]; reviewed in [43]), and across gametes produced by the same individual [36, 38, 44–48]. Finally, comparisons have been made of the levels of shuffling within different chromosomes [49–51], with implications for which chromosomes are most susceptible to harboring selfish genetic complexes [12] or to be used as new sex chromosomes [25, 52].

Quantitative comparisons of such types require a proper measure of genome-wide shuffling. Shuffling is caused both by crossing over and by independent assortment of homologous chromosomes,^1^ which comprise ‘intra-chromosomal’ and ‘inter-chromosomal’ shuffling, respectively.^2^ In previous studies, the most widely used measures of shuffling have considered only the contribution of crossing over, and, more specifically, total crossover frequency or map length. Crossover frequency is simply the number of crossovers that occur during meiotic prophase, as measured either cytologically or from sequence data (further discussion below). Map length is average crossover frequency multiplied by 50 centiMorgans (cM), 1 cM being the map distance between two linked loci that are shuffled in 1% of gametes [53]. Another measure that is sometimes used is the number of crossovers in excess of the haploid chromosome number [14]. Since each bivalent usually requires at least one crossover for its chromosomes to segregate properly [54], the ‘excess crossover frequency’ is the number of crossovers that contribute to shuffling beyond this supposed structurally-required minimum. None of the above measures takes into account shuffling caused by independent assortment of homologs. A fourth measure of aggregate shuffling, which does take into account independent assortment, is Darlington’s ‘recombination index’ (RI) [1, 55, 56], defined as the sum of the haploid chromosome number and the average crossover count. The rationale for this measure derives from the fact that, given no chromatid interference in meiosis [53, 57], two linked loci separated by one or more crossovers shuffle their alleles with probability 1/2 in the formation of a gamete, as if the loci were on separate chromosomes. The RI is therefore the average number of ‘freely recombining’ segments per meiosis.

Importantly, none of these existing measures takes into account the specific positions of crossovers on the chromosomes. Intuitively, though, crossover position is a critical parameter. For example, a crossover at the far tip of a pair of homologous chromosomes does little work in shuffling the genetic material of those chromosomes, while a crossover in the middle causes much shuffling (Fig. 1). Additionally, two crossovers close together may cancel each other’s effect (except for the few loci that lie between them) and thus result in less allelic shuffling than two crossovers spaced further apart.

**Figure 1:**
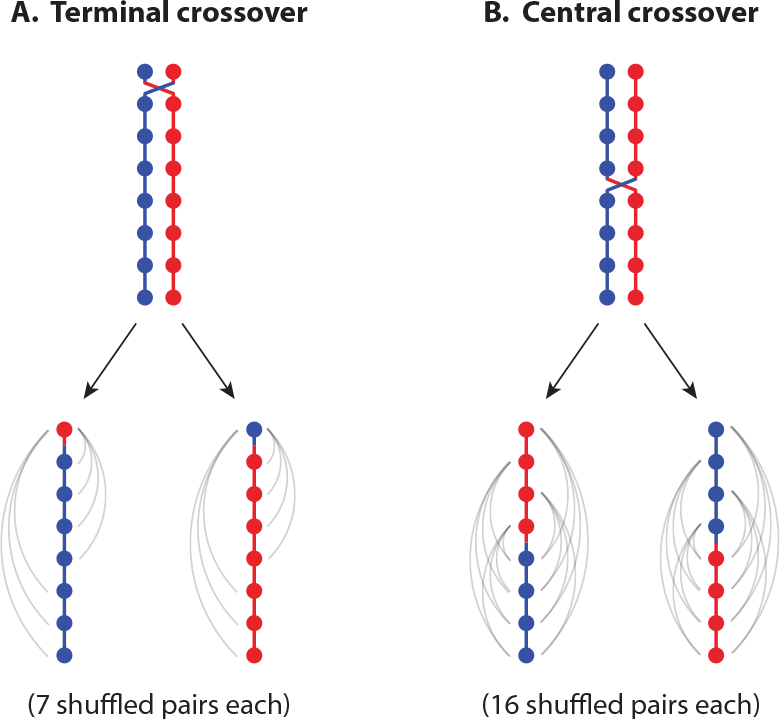
The position of a crossover affects the amount of genetic shuffling that it causes. The figure shows the number of shuffled locus pairs that result from crossing over between two chromatids in a one-step meiosis. The chromosome is arbitrarily divided into eight loci. (A) A crossover at the tip of the chromosome, between the seventh and eighth loci, causes 7 of the 28 locus pairs to be shuffled in each resulting gamete. (B) A crossover in the middle of the chromosome, between the fourth and fifth loci, causes 16 of the 28 locus pairs to be shuffled in each resulting gamete. The central crossover thus causes more shuffling than the terminal crossover.

Therefore, existing measures do not actually define the total genome-wide amount of shuffling, instead serving only as proxies for this critical parameter. This is not a trivial concern. There is significant heterogeneity in the chromosomal positioning of crossovers at all levels of comparison—between species [15, 58, 59], populations within species [60–64], the sexes [13, 25, 65–68], individuals [41, 69–72], and different chromosomes [49, 68, 73, 74], all of which will impact the level of shuffling that these crossovers will cause. Moreover, there can be major differences in chromosome number and size across species [1, 75, 76], which will seriously influence the total amount of shuffling due to independent assortment.

The fact that the positions of crossovers matter for the total amount of shuffling has been recognized for many years (e.g., [1, 65, 75, 77–79]). The need for a measure of total shuffling that accounts for crossover positions has also previously been recognized. Indeed, Burt, Bell, and Harvey [26] give an explicit formula, identical to (3) below, for ‘the proportion of the genome which recombines’ (in the population genetics sense—see footnote 2). Colombo [80] also gives an explicit characterization of what we call *r̅*, phrased as a generalization of the RI. Finally, Haag et al. [81], after noting that terminal crossovers cause little shuffling and that map length is therefore an imperfect measure of genome-wide shuffling, suggest a better measure to be ‘the average likelihood that a [crossover] occurs between two randomly chosen genes’. However, no mathematical expression that incorporates crossover position in the measurement of genome-wide shuffling has been developed and implemented.

Here, we present a simple, intuitive measure of the genome-wide level of shuffling. We define *r̅* as the probability that a randomly chosen pair of loci shuffle their alleles in meiosis, taking into account crossover number and position and the contribution of independent assortment. We have chosen the name ‘*r̅*’ to echo classic population genetics terminology, where the parameter ‘*r*’ for a given pair of loci is the probability that they shuffle their alleles in a gamete. Our parameter *r̅* is simply this quantity averaged across all locus pairs.

In the present work, we define the quantity *r̅*, develop it mathematically and statistically, and document its intrinsic implications, e.g., for crossover interference. We also show how it can be decomposed into separate components deriving from crossovers and from independent assortment of homologs. We then discuss the approaches now available to allow chromosome-specific and/or genome-wide measurement of crossover positions. With these developments in hand, we present a first application of *r̅* to quantitative evaluation of shuffling in human males and females.

## Derivation of *r̅*

*r̅* is the probability that the alleles at two randomly chosen loci will be shuffled in the production of a gamete. It can be calculated with knowledge of crossover positions on each chromosome, in conjunction with knowledge of the fraction of the total genomic length (in bp) accounted for by each chromosome (chromosome lengths influence shuffling caused by both independent assortment and crossovers). In what follows, we always assume that there are sufficiently many loci that the difference between sampling them with or without replacement is negligible.

### Formulas for *r̅*

In the ideal situation, crossover positions are defined for individual gametes, and the parental origins of the chromosome segments delimited by these crossovers are known. In this case, the proportion *p* of the gamete’s genome that is paternal can be determined, and the probability that the alleles of a randomly chosen pair of loci were shuffled during formation of the gamete, either by crossing over or by independent assortment, is simply the product of the probabilities that one locus is of paternal origin (probability *p*) and the other of maternal origin (probability 1 − *p*), in each of the two possible combinations:

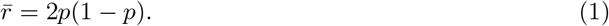

Such data can emerge directly from sequencing of single (haploid) gametes (Fig. 2B) or of a diploid offspring in which the haploid contribution from a single gamete can be identified (Fig. 2C).

**Figure 2:**
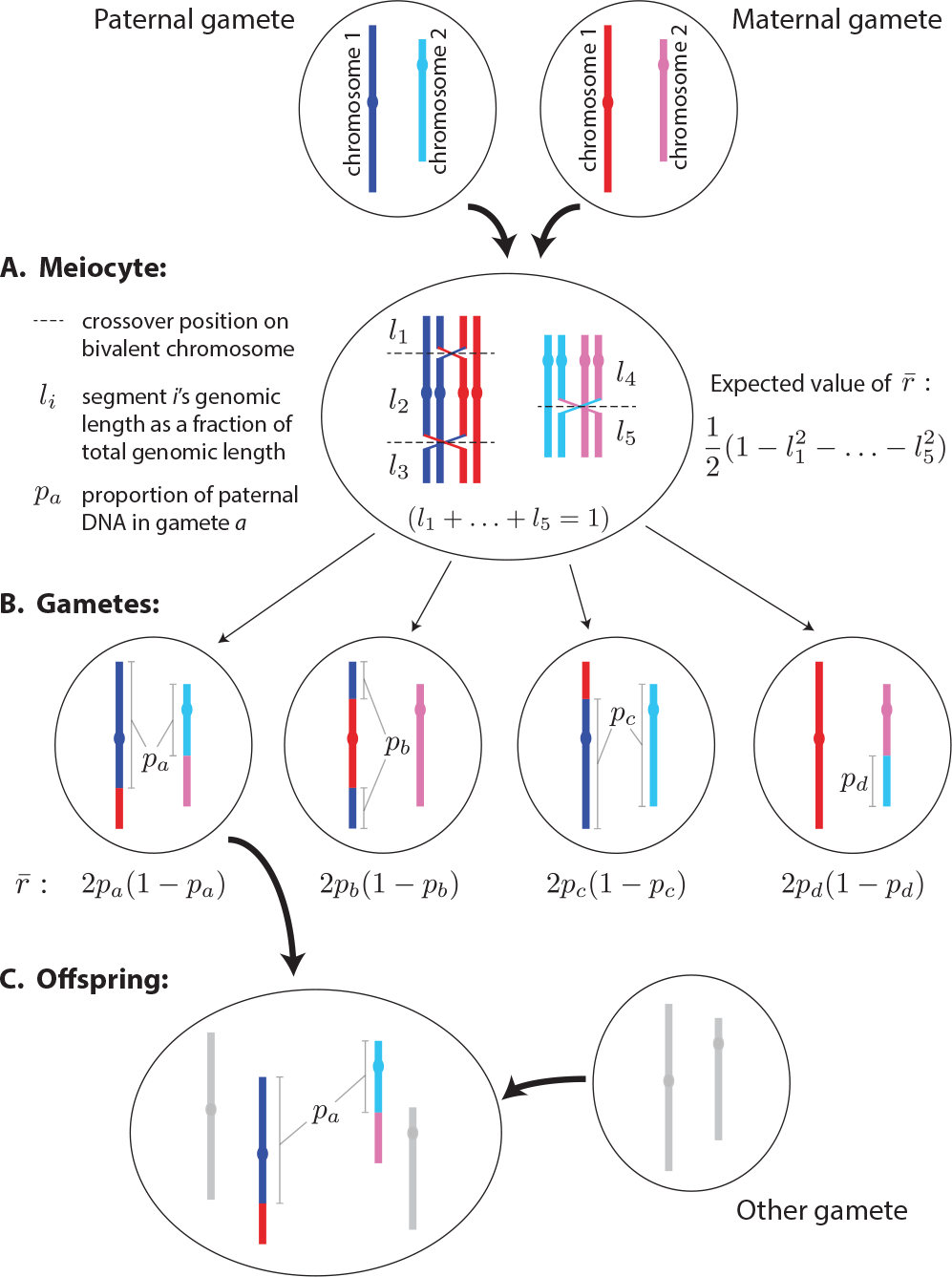
(A) Calculating the expected value of *r̅*, 𝔼[*r̅*], at meiosis I using (3). (B,C)Calculating *r̅* in resulting gametes or offspring using (1).

In some analyses, while crossover positions for a gamete are known, the specific parental identities of the chromosome segments that these crossovers delimit are not known, because the gamete-producer has been sequenced but its parents have not. In this case, we can nonetheless calculate an expected value of *r̅* for the gamete. The parental origin of segments will alternate across crossovers on each chromosome in the gamete, and so we can still define, for any given chromosome *k*, the proportions that originate from the two different parents (*p*_*k*_ and 1 − *p*_*k*_). However, it is not possible to know, from one chromosome to the next, which sets of segments are from the same or different parents. We address this ambiguity by assuming that the probability that the alleles at two loci on separate chromosomes have been shuffled is 1/2, consistent with Mendel’s Second Law. If there are *n* chromosomes in the haploid set, and chromosome *k* accounts for a fraction *L*_*k*_ of the genome’s length, and we determine that a proportion of chromosome *k* is of one parental origin and 1 − *p*_*k*_ is of the other, then

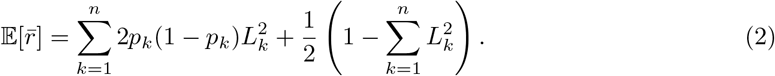

The first term in (2) is the probability, summed over all chromosomes *k*, that two randomly chosen loci lie on chromosome *k* multiplied by the probability that, if so, they are of different parental origin. The second term is the probability that the two randomly chosen loci lie on different chromosomes, 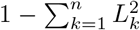 (i.e., one minus the probability that a random pair of loci lie on the same segment), multiplied by 1/2. This second term is probabilistic, and so (2) is an expectation of *r̅* rather than the value actually realized in the gamete.^3^

Finally, cytological analysis of meiotic pachytene chromosomes allows to define crossover positions along bivalent chromosomes at that stage of meiotic prophase I. An expected value for *r̅* can be calculated in this case following the formulas given by Burt, Bell, and Harvey [26] and Colombo [80] (Fig. 2A). If the haploid number of chromosomes is *n* and there are a total of *I* crossovers, then these crossovers divide the bivalents into *n* + *I* segments (this is the RI described above). Label the segments in some (arbitrary) order, and suppose that segment *i*’s fraction of total genome length is *l*_*i*_, with *l*_1_ + … + *l*_*n*+*I*_ = 1. For any randomly-selected locus pair to shuffle its alleles in the production of a gamete, the two loci need to be situated on different segments, the probability of which is 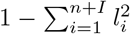. If the two loci are indeed situated on different segments, they shuffle their alleles with probability 1/2 (this assumes no chromatid interference, so that two linked loci separated by one or more crossovers at meiosis I shuffle their alleles in a resulting gamete with probability 1/2 [53, 56]). Therefore, given the configuration of crossovers at meiosis I, the probability that the alleles at a randomly chosen pair of loci will be shuffled in a resulting gamete is 1/2 multiplied by the probability that the two loci are on separate segments, or

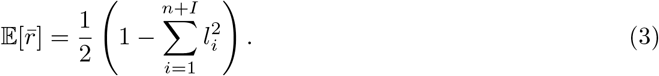

𝔼[*r̅*], as defined by (3), is proportional to the Gini-Simpson index [82], a commonly used measure of diversity, especially in ecology [83].

### Inter- and intra-chromosomal components of *r̅*

It is often argued that the predominant source of shuffling in sexual species is independent assortment of separate chromosomes, or, correspondingly, that the most effective way for a species to increase genome-wide shuffling in the long term is to increase the number of chromosomes rather than crossover frequency ([1, 84]; though see [85] and *Discussion*). Our formulation of *r̅* allows us to partition total shuffling into a component deriving from crossovers (intra-chromosomal shuffling) and a component deriving from independent assortment of separate chromosomes (inter-chromosomal shuffling), thereby allowing us to evaluate previous assertions about the relative contributions of each in a rigorous, quantitative way.

The inter-chromosomal component of *r̅* is the probability that two randomly chosen loci are on separate chromosomes and shuffle their alleles, while the intra-chromosomal component is the probability that two loci are on the same chromosome and shuffle their alleles. We first present this decomposition for *r̅* in a gamete or haploid complement of an offspring. If chromosome *k* contains a proportion *p*_*k*_ of paternal content, then the appropriate decomposition is

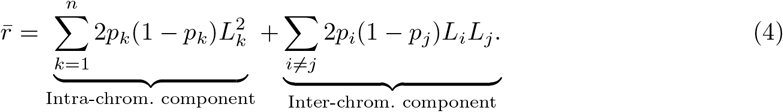

If we know crossover realizations in a haploid complement but not the specific parental origin of the segments they delimit, then we can still calculate a proportion *p*_*k*_ of chromosome *k* to be of one parental origin and 1 − *p*_*k*_ of the other (though not knowing which is which), so we retain the first term in (4) but lose information about the second term. The appropriate partition is then

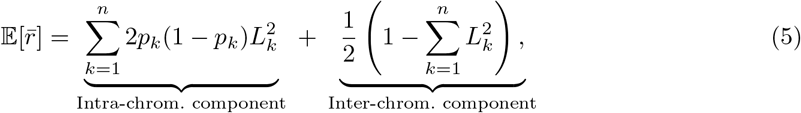

which is the same as (2).

We now present the decomposition for 𝔼[*r̅*] at meiosis I. Suppose that the haploid number of chromosomes is *n*, and that bivalent *k* exhibits crossovers, dividing it into segments *i* = 1,…, *I*_*k*_ + 1, whose fractions of the total genomic length are *l*_*k,i*_. Chromosome *k*’s fraction of the total genomic length is 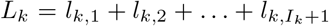, with *L*_1_ + *L*_2_ + … + *L*_*n*_ = 1. Then (3) is partitioned as follows:

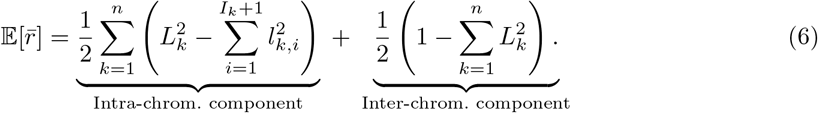

### Averaging *r̅*

The above measures can be aggregated to obtain the average value of *r̅* across many gametes (or meiocytes). We denote the average value of a variable *x* by 〈*x*〉. To average *r̅* given sequence data from many gametes, and supposing that, for each gamete, we can distinguish which sequences are paternal and which are maternal, we can take the average value of (4), noting that the *L*_*k*_ are constants:

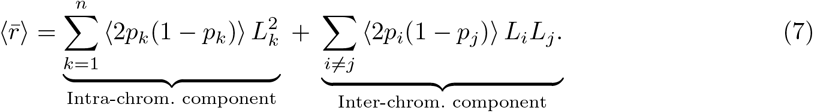

If segregation is Mendelian, then in a large sample of gametes, 〈2*p*_*i*_(1 − *p*_*j*_)〉 ≈ 1/2, and so the inter-chromosomal component in (7) will be close to 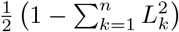.

If we have sequence data from many gametes and can determine crossover positions but not the parental origin of the sequences these crossovers delimit, then we take the average of (5), noting again that the *L*_*k*_ are constants (which, here, means that the inter-chromosomal component is constant):

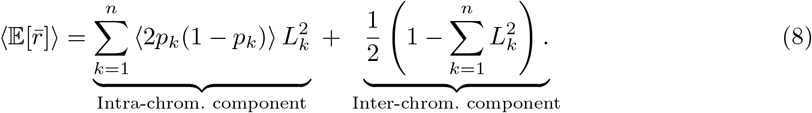

Given data of crossover positions along bivalent chromosomes in many meiocytes, we take the average of (6):

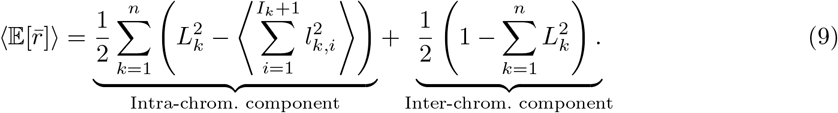

(7)-(9) mean that we can estimate the average intra-chromosomal contribution to *r̅* separately for each chromosome, possibly from different sets of data, and then combine these averages into a final measure of *r̅*. This is useful, because often in sequencing or cytological studies of large numbers of gametes or meiosis I nuclei, it is possible (or desired) to obtain accurate measurements for only a subset of the chromosomes in each cell, so that the sets of cells from which measurements are taken for two chromosomes will in general not overlap. This is the case, for example, in the very rich cytological data from human pachytene nuclei in [47]. In such cases, (7)-(9) state that the average intra-chromosomal contribution of one chromosome can be estimated from the set of cells from which measurements of that chromosome were possible, while the contribution of another chromosome can be estimated from the set of cells from which measurements of that chromosome were possible (possibly from different sets of data); these two separate estimates can then be combined in (7)-(9).

Unrelated to the averaging of *r̅* across multiple measurements of individual gametes or meiocytes, an average value for *r̅* can also be calculated directly given pairwise average rates of shuffling for all loci. As discussed below, these can be estimated from linkage maps generated from pooled sequence data. Suppose that we have measured, for each locus pair (*i, j*), their rate of shuffling *r*_*ij*_. 〈*r̅*〉 is then simply the average value of *r*_*ij*_ across all locus pairs (*i, j*). If Λ is the total number of loci, then

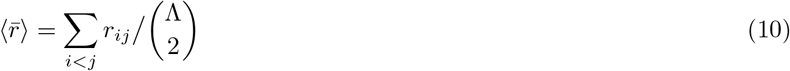

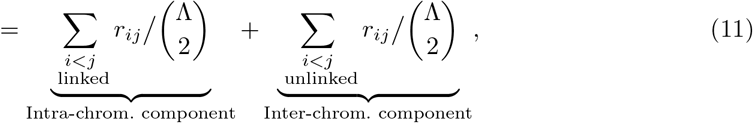

where 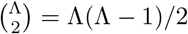. When *r*_*ij*_ = 1/2 for all unlinked locus pairs (*i,j*), the inter-chromosomal component simplifies to 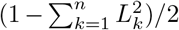, where *L*_*k*_ is chromosome *k*’s proportion of total genomic length.

## Properties and intrinsic implications of *r̅*

### Properties

We note three properties of *r̅*. First, its minimum value is 0, and its maximum value is 1/2, the latter relying on our assumption of many loci. The maximum value of *r̅* in a gamete can occur by chance equal segregation of maternal and paternal DNA to the gamete. The maximum value of 𝔼[*r̅*] at meiosis I requires, unrealistically, that every pair of loci are either on separate chromosomes or experience at least one crossover between them in every meiosis. The minimum value of *r̅* in a gamete could result from chance segregation of only crossover-less chromatids of one parental origin to the gamete. The minimum value of 𝔼[*r̅*] at meiosis I requires crossing over to be absent (as in *Drosophila* males and *Lepidoptera* females [75]), and either a karyotype of one chromosome or a meiotic process that causes multiple chromosomes to segregate as a single linkage group (as in some species in the evening primrose genus, *Oenothera* [86]).

Second, *r̅* satisfies the intuitive property that a crossover at the tip of a chromosome causes less shuffling than a crossover in the middle (some example calculations of *r̅* are given in Fig. 3). This is easily seen in (3). Consider a bivalent chromosome on which a single crossover will be placed. This will divide the chromosome into two segments of length *l*_1_ and *l*_2_, where *l*_1_ + *l*_2_ = *L* is the chromosome’s fraction of total genomic length. The contribution of this crossover to 𝔼[*r̅*] is seen from (3) to be 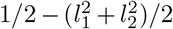, which can be rewritten *l*_1_*l*_2_ + 1/2 − (*l*_1_ +*l*_2_)^2^/2, which simplifies to *l*_1_*l*_2_ under the constraint *l*_1_ +*l*_2_ = *L*. *l*_1_*l*_2_ is maximized when *l*_1_ = *l*_2_ = *L*/2, i.e., when the crossover is placed in the middle of the bivalent. The quantity is minimized when either *l*_1_ = 0 or *l*_2_ = 0, i.e., when the crossover is placed at one of the far ends of the bivalent. In general, as the crossover is moved further from the middle of the bivalent, 𝔼[*r̅*] steadily decreases. This is true regardless of the positions of the crossovers on other chromosomes. Importantly, this relationship also holds if, instead of considering where to place a single crossover on a whole bivalent, we consider where to place a new crossover on a segment already delimited by two crossovers (or by a crossover and a chromosome end). In general, if we are to place any fixed number of crossovers along a single chromosome, 𝔼[*r̅*] is increased if they are evenly spaced (SI Appendix).

**Figure 3:**
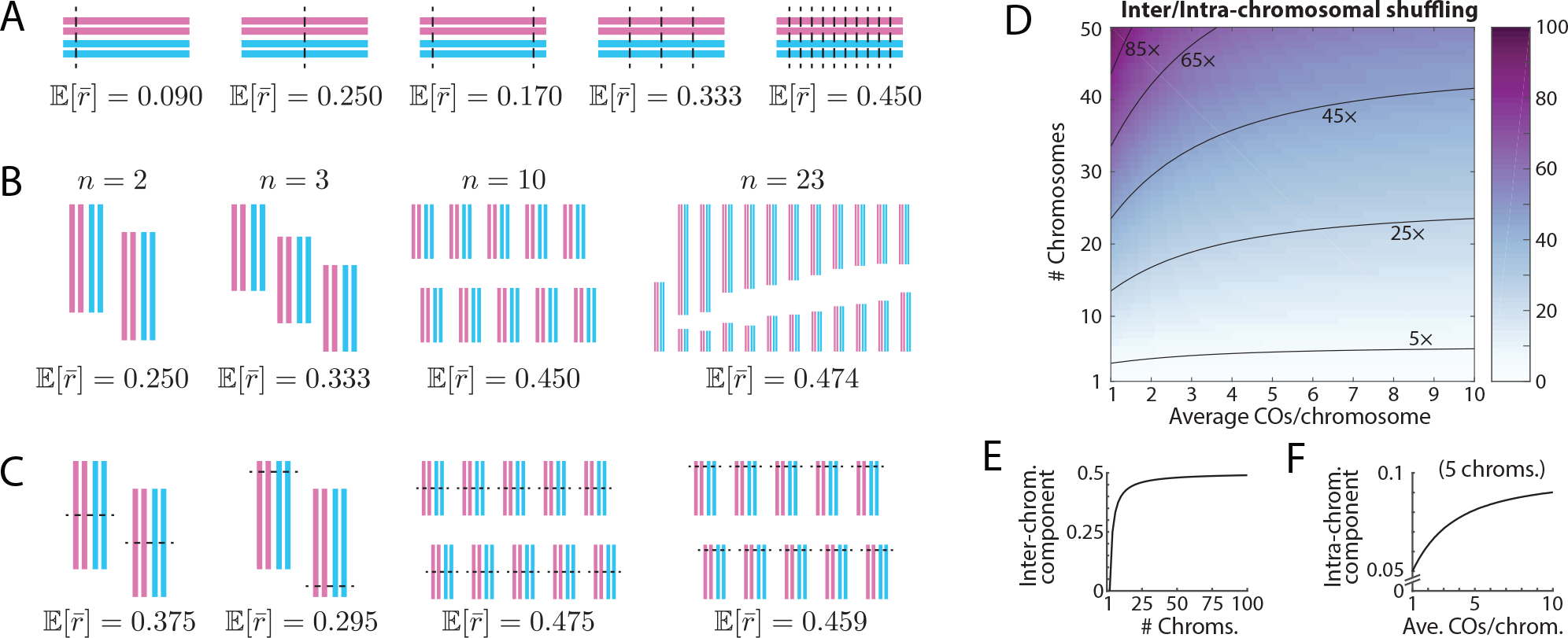
(A-C) Example values of the expected value of *r̅*, 𝔼[*r̅*], at meiosis I. In each case, 𝔼[*r̅*] is calculated according to (3). (A) Various crossover numbers and positions in a diploid organism with only one chromosome. Notice that 𝔼[*r̅*] increases sub-linearly with crossover number, even when crossovers are evenly spaced so as to maximize 𝔼[*r̅*]. (B) Various chromosome numbers for a diploid organism without crossing over. Notice that 𝔼[*r̅*] increases sub-linearly with chromosome number, even when chromosomes are evenly sized so as to maximize 𝔼[*r̅*]. Notice too that, when the number of chromosomes is not small, 𝔼[*r̅*] is close to its maximum value of 1/2. The rightmost panel represents a crossover-less meiosis in a human female, in which case 𝔼[*r̅*] is 0.474, close to its maximum value of 1/2. (C) Various configurations of crossovers in a diploid organism with multiple chromosomes. (D) The ratio of contributions to 𝔼[*r̅*] from independent assortment of homologs (inter-chromosomal shuffling) and crossing over (intra-chromosomal shuffling). Chromosomes are assumed to be of equal size, and crossovers are assumed always to be evenly spaced along chromosomes. For each combination of *n* ≥ 2 chromosomes in the haploid set and *I* ≥ 1 crossovers per chromosome on average, we simulate 10^5^ nuclei. For each of the *n* bivalent chromosomes in a nucleus, we draw a value from a Poisson distribution with parameter *I* − 1 and add it to 1 mandatory crossover to get the total number of crossovers for that chromosome. We then calculate the average total intra-chromosomal contribution to 𝔼[*r̅*] across the nuclei, and compare it to the inter-chromosomal contribution (which is constant across nuclei). When the number of chromosomes is not small, the inter-chromosomal component dominates. (E) The inter-chromosomal component increases at a decreasing rate as chromosome number increases. (F) Similarly, holding constant the number of chromosomes (here, 5), the intra-chromosomal component increases at a decreasing rate as average crossover number increases.

Third, *r̅* will tend to increase sub-linearly with both the number of crossovers and the number of chromosomes (Fig. 3D,E,F). For example, increasing the number of crossovers by some factor will increase the intra-chromosomal component of *r̅* by a lesser factor (and *r̅* as a whole by a yet lesser factor, because its inter-chromosomal component is unaffected). Similarly, doubling the number of chromosomes in the haploid set will cause a less-than-twofold increase in the inter-chromosomal component of *r̅*. That is, there are ‘diminishing returns’ to genome-wide shuffling from having increasingly more crossovers and more chromosomes. These effects are described in mathematical detail in SI Appendix.

### Intrinsic implications

We have noted above that, given some number of crossovers along a chromosome at meiosis I, 𝔼[*r̅*] is maximized if they are evenly spaced. This observation carries the interesting implication that positive crossover interference—a classical phenomenon [87, 88] where the presence of a crossover at some point along a bivalent chromosome inhibits the formation of nearby crossovers—will tend to increase *r̅*. It thus suggests a possible selective advantage for this phenomenon (see Discussion).

Also, the general ‘diminishing returns’ to *r̅* of having more crossovers (above) carries particular implications for differences in total autosomal shuffling in the two sexes of a species. When the number of crossovers and/or their localization does not differ grossly between the sexes, then the amount of shuffling will be similar in both sexes (e.g., for male and female humans, as discussed below).

## Experimental determination of crossover number and position

Measurement of the quantity *r̅* requires that crossover positions along chromosomes can be accurately determined. While this was previously not possible, cytological advances have made it possible to efficiently and accurately visualize the positions of crossovers on pachytene bivalent chromosomes [36, 73, 74, 89, 90], while rapid technological advances in DNA sequencing have allowed crossover positions to be determined at a fine genomic scale using sequence/marker analysis of pedigrees [39, 40, 91, 92], individual gametes [93, 94] and meiotic triads/tetrads [41, 95–97]. These technological advances allow simple estimation of *r̅*.

### Cytological analysis

The physical positions of crossovers along the axes of pachytene chromosomes can be determined reliably in spread pachytene nuclei (or in 3D reconstructions from serial sectioning), either with electron microscopy by direct visualization of ‘late recombination nodules’ (which mark all crossovers [98–101]), or with light microscopy using immunofluorescent staining techniques to detect ‘type I’ (interfering) crossovers (reviewed in [102]; see Fig. 4), which are the vast majority in most organisms [103]. The latter method is now most commonly used.

**Figure 4:**
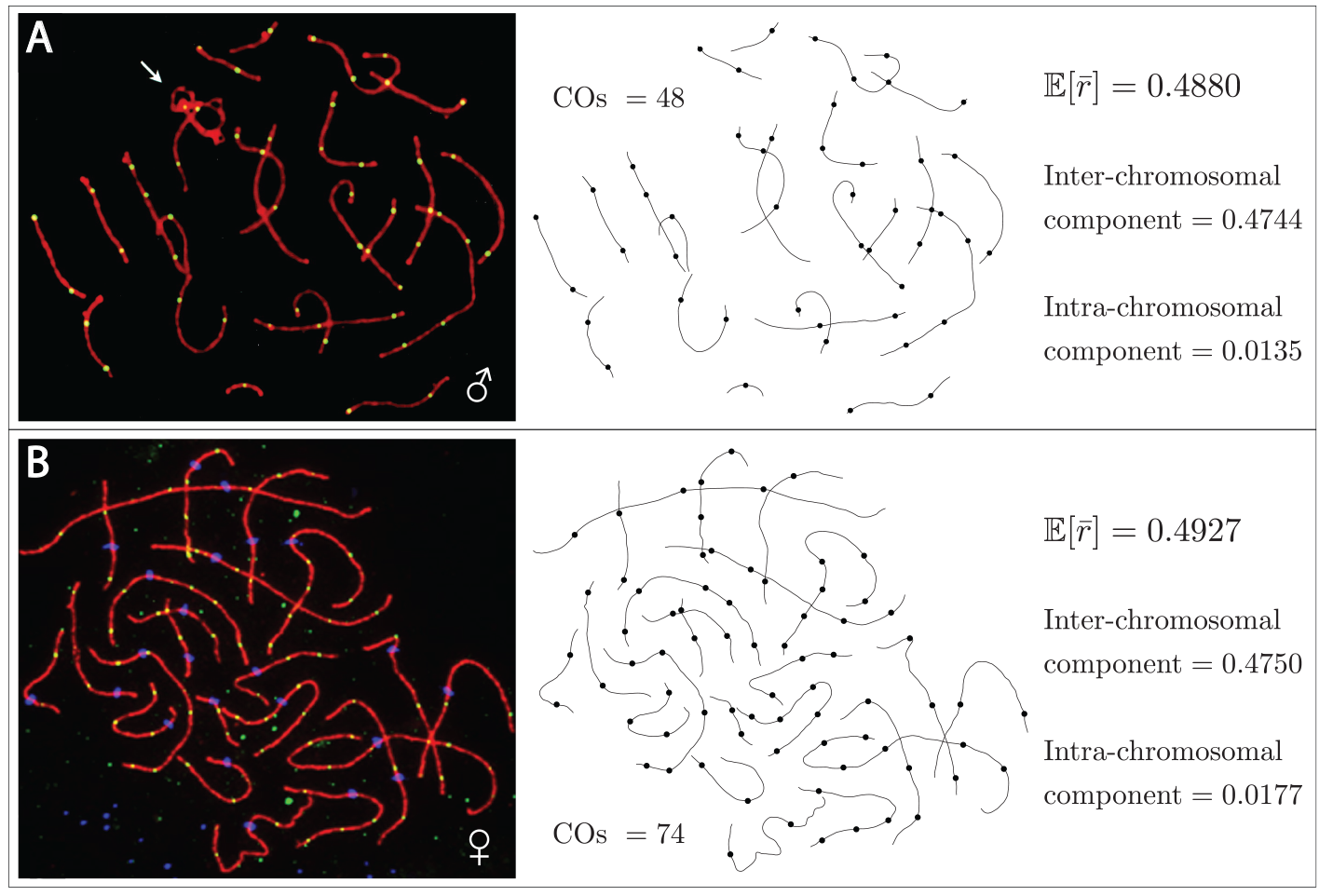
Calculation of 𝔼[*r̅*] from crossover patterns at meiosis I in humans. The left panels are micrographs of immunostained spreads of a pachytene spermatocyte (A) and a pachytene oocyte (B). The red lines show localization of synaptonemal complex protein SYCP3, and demarcate the structural axes of the chromosomes. The green dots along the axes show foci of the protein MLH1, and demarcate the positions of crossover associations along the axes. The white arrow in (A) points to the paired X and Y chromosomes. In (B), the blue dots along the chromosome axes mark the centromeres. The paired X chromosomes in (B) have not been identified, and are included in the calculation of 𝔼[*r̅*]. The central panels show the traced axes and crossover positions from the spreads. From these, the lengths of the segments separated by the crossovers can be measured, and converted to fractions of the total physical axis length (these measurements can be found in Supplement File S1). The segments’ fractions of total physical length can be taken to be approximately equal to their fractions of total genomic length, for reasons described in the text. With these fractions of genomic length, 𝔼[*r̅*] may be calculated for each spread according to (3), and its inter- and intra-chromosomal components determined according to (6). The results of these calculations are given in the right panels. Notice that we do not require individual chromosomes to be identified to calculate 𝔼[*r̅*] or its intra-and inter-chromosomal components. The left panels of (A) and (B) are modified, with permission, from [102] and [38] respectively.

Visualization of the chromosome axes can be achieved by staining either the axes or the SC central region (e.g., SYCP1-3). Crossover positions along the axes can be visualized by staining molecules specific to crossover recombination complexes (e.g., MLH1). If desired, the positions of centromeres along the axes can be visualized by staining for centromeric proteins such as CENP-A, and individual chromosomes can be identified by fluorescence in situ hybridization (FISH) or locus-specific fluorescent tags (e.g., FROS arrays), or, in favorable cases (e.g., *Arabidopsis* [104]), by relative chromosome length. Spread pachytene nuclei immuno-stained for crossover positions are now common in the literature, and have been generated for many species (e.g., 28 bovid species in [105] alone).

Only physical distances (in *μ*m) along chromosomes can be directly inferred from cytological data, and so the measurement of *r̅* from such data requires a way to convert physical distances into genomic distances (in bp). The use of pachytene-stage chromosomes in the measurement of *r̅* (Fig. 4) is particularly advantageous in this regard because, at this stage of meiosis, genomic distance is proportional to physical distance along the chromosome axes (below and SI Appendix).

The physical structure of these chromosomes is highly regular: The chromatin of each chromatid is arranged in a linear array of loops, the bases of which lie at regular intervals along a common axis (reviewed in [100, 106, 107]). Almost all of the DNA is accommodated in the loops, with very little DNA in the axis between loop bases [100, 106–108]. Therefore, the genomic lengths of the loops directly define the genomic distance per unit of physical distance along a pachytene bivalent axis (the local ‘packing ratio’) [100, 105, 107, 109, 110]. Moreover, the local packing ratio along and across different chromosomes is approximately constant within each nucleus, at least at the relatively crude scale of inter-crossover distances required for analysis of *r̅*, as indicated by two criteria. First, physical loop lengths can be measured directly, and, in a variety of species, appear to vary minimally along and across chromosomes within nuclei (e.g., [106, 110–113]; reviewed in [100]). Second, the relative genomic lengths of chromosomes closely match their relative physical axis lengths at pachytene (see Fig. S2 for examples from diverse species), from which we infer that the average packing ratio of different chromosomes within a nucleus is approximately constant. Given these considerations, the chromosome segments’ fractions of total genomic length (bp), as required for calculation of 𝔼[*r̅*], correspond to, and thus can be defined by, their fractions of total physical axis length (*μ*m). Therefore, their fractions of total axis length (easily measured from cytological spreads) can be substituted into the formulas derived above for 𝔼[*r̅*] and its components.^4^

### Sequence analysis

The genomic positions of crossovers in an individual’s meioses can be inferred by sequencing the individual products of those meioses—either directly by sequencing individual gametes or polar bodies, or by inferring gametic genomes by sequencing individual diploid offspring—provided the individual’s diploid genome has been haplophased. Haplophasing can be achieved either by sequencing an extended pedigree involving the individual [39, 72, 115], by sequencing multiple offspring, gametes, and/or polar bodies of the individual [41, 51, 94, 96, 116, 117], or by isolating individual chromosomes from a somatic cell and sequencing them separately [93, 118]. *r̅* can be calculated directly from gametic genome sequencing.

*r̅* can also be calculated from linkage maps derived from pooled sequencing data (as obtained, for example, from population pedigrees [39] or pooled sperm [119]). A linkage map gives the map distance (in cM) between pairs of linked markers. Using this, we can generate an evenly spaced grid of loci for each chromosome (to ensure even sampling of loci in the calculation of *r̅*), and impute map distances between these pseudomarkers by linear interpolation from distances between true markers in the linkage map (details in SI Appendix). Thus, we generate a map distance *d*_*ij*_ between every pair of linked loci (*i, j*), which we can convert to a rate of shuffling *r*_*ij*_ using a map function [53], e.g., Kosambi’s [120]: 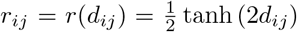. This gives a rate of shuffling for every linked locus pair, from which we can calculate the average intra-chromosomal component of 〈*r̅*〉 using (11). Assuming that unlinked locus pairs shuffle their alleles with probability 1/2, we can calculate the average value of shuffling across all locus pairs, 〈*r̅*〉, using (10).

### Using simulation data to estimate *r̅*

One potential drawback of measuring *r̅* is that the data required, such as cytological spreads and gametic genome sequences as described above, can be laborious or expensive to obtain in large quantities. This can limit sample sizes, especially in non-model organisms. The beam-film model [121] is a physical model of crossing over that, when computationally calibrated [122], has been shown to accurately reproduce crossover distributions in various taxa and across the sexes [48, 114]. The model can be calibrated given limited input data, after which large data sets of crossovers can be simulated; from these, *r̅* and its components can be calculated. Tweaking various parameters in the model, such as the strength of crossover interference, would also allow study of how these parameters influence *r̅*.

## Measuring *r̅* in humans

### Analysis of cytological data

Fig. 4 shows the calculation of 𝔼[*r̅*] from the immunostained pachytene chromosomes of a single human spermatocyte (from [102]) and oocyte (from [38]). For both spermatocyte and oocyte, 𝔼[*r̅*] is close to its maximum value of 1/2. This high level of shuffling is due almost entirely to independent assortment: in both cases, the inter-chromosomal component of 𝔼[*r̅*] is much larger than the intra-chromosomal component (from crossovers), by a factor of about 35 in the male and about 27 in the female.

Owing to the male-heterogametic (XX/XY) system of sex determination in humans, the spermatocyte in Fig. 4A contains easily identified, partially paired X and Y chromosomes, while the oocyte in Fig. 4B contains a bivalent of paired X chromosomes which in this case has not been distinguished from the other bivalents (e.g., by FISH). Therefore, the X chromosome is included in the calculation for the oocyte, and so the inter-chromosomal components of 𝔼[*r̅*] for the spermatocyte and oocyte are not expected to be equal. Though we have estimated them directly from the single cytological spreads in Fig. 4, they can be calculated exactly from known genomic lengths of the chromosomes. Substituting the chromosome lengths reported in assembly GRCh38.p11 of the human reference genome (available online at www.ncbi.nlm.nih.gov/grc/human/data?asm=GRCh38.p11; lengths listed in Supplement File S1) into the first term in (6), we find that the value for the spermatocyte should be 0.4730, close to the value of 0.4744 calculated in Fig. 4A, while the value for the oocyte should be 0.4744, close to the value of 0.4750 calculated in Fig. 4B. This close match between the exact values and those deduced from physical chromosome axis lengths illustrates the reliability of physical length as a proxy for genomic length in the calculation of 𝔼[*r̅*].

Interestingly, although the female spread exhibits about 50% more crossovers than the male spread (74 vs. 48), the intra-chromosomal component of 𝔼[*r̅*] for the female spread is only about 30% larger than that of the male spread. This is expected from the general arguments given earlier concerning the ‘diminishing returns’ of more crossovers (Fig. 3D,F).

We have carried out measurement of 𝔼[*r̅*] using data from 824 spermatocytes collected from 10 male humans (data from [123], kindly provided by F. Sun). Using (9), we calculate each chromosome’s average contribution to the intra-chromosomal component of 𝔼[*r̅*] (these are listed in SI Table S1). Summing these values, we find that the average intra-chromosomal component of 𝔼[*r̅*] is 0.0143, which is similar to the value calculated for the single spermatocyte in Fig. 4A. Restricting attention to the 715 cells in which data were collected for every chromosome allows us to calculate the average and standard deviation of the intra-chromosomal component of 𝔼[*r̅*] for each of the 10 individuals. The standard deviation of the intra-chromosomal component of 𝔼[*r̅*] across the spermatocytes of each individual is of order 0.0015 (SI Table S2), or about 10% of the average value. There is some variation among the individuals for the average value of the intra-chromosomal component of 𝔼[*r̅*]: the minimum value is 0.0130, while the maximum is 0.0153. Interestingly, these minimum and maximum average values do not come from the individuals with, respectively, the smallest and largest average crossover frequencies (SI Table S2). In fact, the individual with the largest average value of the intra-chromosomal component of 𝔼[*r̅*] has only the fourth (out of ten) largest average number of crossovers per spermatocyte. This further highlights the importance of crossover position for the amount of genetic shuffling, and hints that individuals might differ systematically in their crossover positioning in ways that quantitatively affect the amount of shuffling in their gametes.

### Analysis of gamete sequence data

Hou et al. [96] sequenced the products of multiple meioses (first polar bodies, second polar bodies, and ova) of several females, and haplophased these females directly from the sequences of the meiotic products, allowing the crossover points in those products to be determined. Fig. 5 is a modified version of Fig. S2 from [96], showing the crossover points along the chromosomes of an egg cell from one of the individuals. Because relatives of the individual were not sequenced, the paternal and maternal sequences cannot be discerned. Therefore, calculation of 𝔼[*r̅*] proceeds according to (2), with its inter- and intra-chromosomal components determined according to (5). Carrying out this calculation for all chromosomes, including the X, reveals a value of 𝔼[*r̅*] = 0.4971, with an inter-chromosomal component of 0.4756 (calculated using chromosome lengths as they appear in Fig. 5) and an intra-chromosomal component of 0.0215 (all details in Supplement File S2). The inter-chromosomal contribution is about 22 times larger than the intra-chromosomal contribution. If we instead restrict attention to the autosomes, we find 𝔼[*r̅*] = 0.4965, with an inter-chromosomal component of 0.4735, and an intra-chromosomal component of 0.0230.

**Figure 5:**
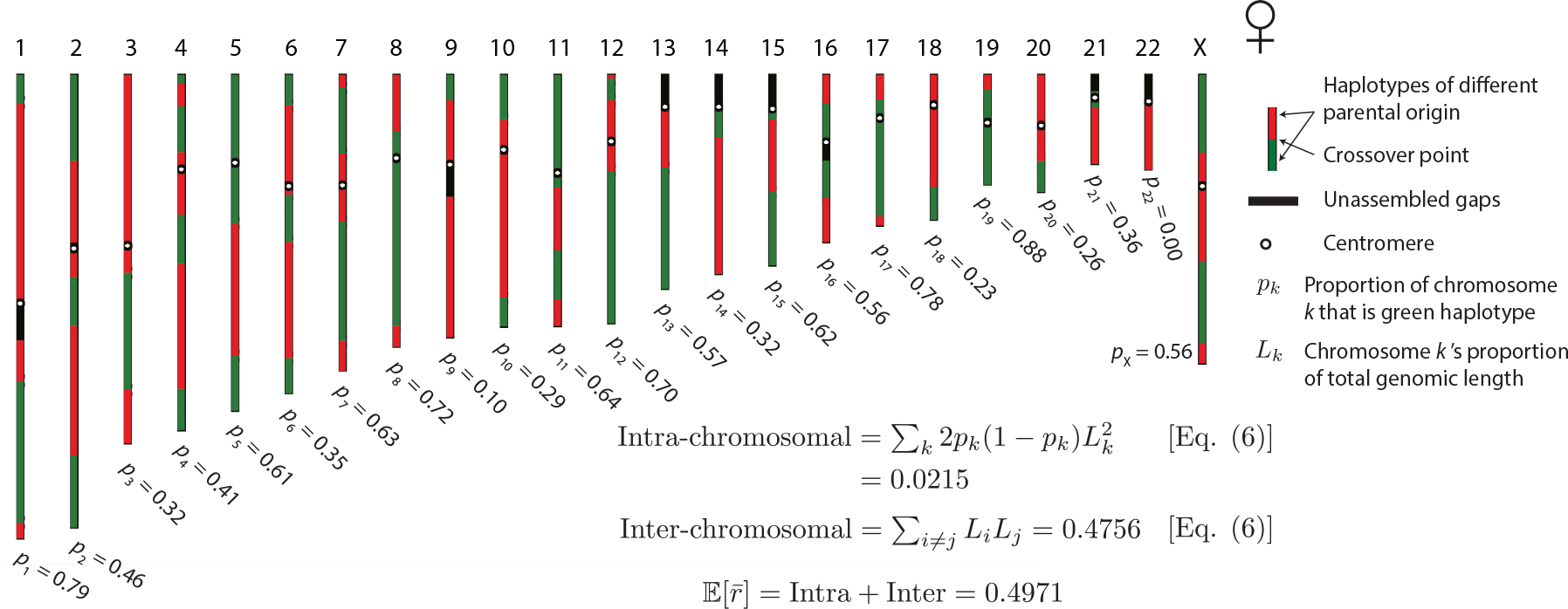
Calculation of *r̅* from the sequence of an individual egg obtained from a haplophased human female. Points of crossing over along chromosomes can be estimated directly from the gamete sequence. With no sequencing of an extended pedigree, we cannot tell which of the haplotypes separated by crossover points are maternal and which are paternal. Therefore, we cannot directly calculate *p*, the proportion of the egg’s DNA that is paternal. We can, however, for each chromosome *k* find the proportion *p*_*k*_ that is of one parental origin (without knowing which), and then use (5) to calculate *r̅* and its components. Modified, with permission, from [96].

Fan et al. [118] haplophased a male using microfluidic techniques, and Wang et al. [93] sequenced 91 individual sperm cells from this male, using the haplophasing from [118] to determine the points of crossing over in each sperm’s sequence (given in Table S2 of [93]). From these, and because maternal and paternal sequences are not known, we use (8) to calculate the average value, across all 91 gametes, of 𝔼[*r̅*] and its components. Carrying out this calculation yields an average value of 𝔼[*r̅*] of 0.4856, with an average inter-chromosomal component of 0.4729, and an average intra-chromosomal component of 0.0127 (MATLAB code for these and the following calculations is provided in Supplement File S3). The average inter-chromosomal contribution is therefore about 37 times larger than the average intra-chromosomal contribution. In calculating the inter-chromosomal component of 𝔼[*r̅*], because some shuffled SNP positions for this individual exceeded the chromosome lengths of the human reference genome, we used the position of the last SNP on each chromosome (as reported in [118] in their Supplemental Data Set 2) as our proxy of that chromosome’s genomic length. Because there is no information about the parental origin of the sequences in each sperm, the inter-chromosomal component for each sperm is an average, deduced from the fixed genomic lengths of the individual’s chromosomes [using (5)]. There is therefore no variance across gametes in this component. The standard deviation of the intra-chromosomal component of *r̅* across gametes is 0.0026, resulting in a coefficient of variation of about 0.2. The minimum value of the intra-chromosomal component across all 91 sperm is 0.0058; the maximum value is 0.0178. The average intra-chromosomal component of this individual is low relative to the average value of the intra-chromosomal component calculated from spermatocytes in the previous subsection (0.0143). This is expected, as he is homozygous for a variant of the *RNF212* gene that, in homozygous form, is associated with a reduction of ~5-7.5% in crossover numbers in males [117, 124]. Indeed, the 91 sperm show, on average, 22.8 crossovers each (minimum 12; maximum 32), as compared to the previously cited averages of 25.9 for Icelandic males [51] and 26.2 for Hutterite males [116].

### Analysis of linkage map data

Kong et al. [40] sequenced 8,850 mother-offspring and 6,407 father-offspring pairs of Icelandics to generate high-density linkage maps for both sexes (data publicly available at http://www.decode.com/addendum). For each sex, we apply the method described above for generating a linkage map of evenly spaced ‘pseudomarker’ loci, and we estimate rates of shuffling for all linked locus pairs from the map distances between them using the Kosambi map function. We then estimate the intra-chromosomal component of 〈*r̅*〉 using (11) (the inter-chromosomal component is calculated form known chromosome lengths). This yields a male value of 0.0138, similar to the value of 0.0143 calculated above from 824 spermatocytes (the chromosome-specific contributions are similar too; SI Table S4). This concordance indicates the validity of the linkage-data approach to estimating *r̅*. The female value, including the X chromosome, is 0.0183, similar to the value of 0.0177 calculated from the single oocyte spread in Fig., but smaller than the value of 0.0215 calculated from the single egg in Fig. 5. Excluding the X chromosome, the female value is 0.0193.

## Discussion

The most common current measures of genome-wide genetic shuffling—crossover frequency and map length—do not take into account the positions of crossovers. This is a significant limitation, because terminal crossovers cause less shuffling than central crossovers. Traditional measures of aggregate shuffling also do not take into account independent assortment of homologs in meiosis. Here, we have proposed a measure of genome-wide shuffling that naturally takes both of these features into account. This measure, *r̅*, is the probability that a randomly chosen pair of loci shuffle their alleles in meiosis.

### *r̅* should be the measure of choice for evaluation and comparison of genetic shuffling

*r̅* is the most direct measure of the aggregate amount of shuffling that occurs in the production of gametes. Application of this measure should therefore significantly improve the possibility of understanding the nature, basis, and significance of genetic shuffling and its variation across different situations (e.g., across species). We hope that this will become common practice where possible. In support of this possibility, we have demonstrated that *r̅* can readily be measured from standard sequence and cytological data.

### Distinguishing the contributions of crossing over and Mendel’s Second Law

We have shown that *r̅* can be decomposed into a component from crossovers and a component from independent assortment of homologs. The relative importance of crossing over and independent assortment as sources of shuffling has long been discussed [1, 14, 84, 85]. Theories of the adaptive value of shuffling fall roughly into two categories: those positing a short-term advantage (offspring diversification; e.g., the Tangled Bank [1, 125] and sib-competition [126, 127] theories) and those positing a long-term advantage (e.g., the ‘Fisher-Muller’ theory that gradually reducing allelic correlations over time allows natural selection to operate more efficiently on genes at different loci [2–6]).

Burt [85] has argued that crossovers are more effective than independent assortment in gradually reducing allelic associations (because independent assortment shuffles genomes at fixed points, whereas crossovers can occur at many points along a chromosome). Under this view, the crossover component of rr would be more important than the independent assortment component for longterm theories of the adaptive value of shuffling. On the other hand, the short-term theories rely on aggregate shuffling *per se*. We have shown, using humans as an example, that independent assortment will typically be the greatest contributor to aggregate shuffling. Therefore, the independent assortment component of *r̅* may be more important than the crossover component for short-term theories of the adaptive purpose of shuffling. Our decomposition of *r̅* into these separate components allows these distinctions to be tested quantitatively.

### Crossover interference increases genetic shuffling

A property of the intra-chromosomal component of *r̅* is that it increases as crossovers become more evenly spread out. Thus, intriguingly, interference among crossovers will tend to increase the amount of shuffling that they cause (similar points have been made by Goldstein, Bergman, and Feldman [128], and Gorlov and Gorlova [129]). In SI Appendix, we quantify this effect for human males using the spermatocyte data from [123], described above. For each chromosome, we calculate an ‘interference-less’ average contribution to intra-chromosomal 𝔼[*r̅*] by resampling, in each spermatocyte, the positions of the crossovers independently from the empirical distribution pooled across all spermatocytes. We find that the observed average value of the intra-chromosomal contribution to 𝔼[*r̅*] is about 15% higher than this interference-less value.

Crossover interference, first described more than a century ago [87], is a deeply conserved feature of the meiotic program [130, 131]. Despite this, its adaptive function remains unclear [103, 132, 133]. One category of hypotheses invokes mechanical advantages of spread-out crossovers, either to bivalent stability in prophase [134] or to homolog segregation at meiosis I [135]. However, such ideas are challenged by the fact that meiotic segregation proceeds without trouble in organisms that lack crossover interference (e.g., fission yeast and *Aspergillus* [136]) or that have had interference experimentally reduced or eliminated [137, 138]. The present study raises the possibility of a qualitatively different idea, that crossover interference provides an evolutionary advantage via its effects on genetic transmission. By explicitly taking into account the positions of crossovers in quantifying how much shuffling they cause, we show that crossover interference serves to increase the shuffling caused by a given number of crossovers. This finding therefore suggests a new possibility for the adaptive function of crossover interference.

### Additional threshold-based measures of genetic shuffling

*r̅* is a suitable measure of the total amount of shuffling, linearly aggregating the rates of shuffling between different locus pairs. In this way, it quantifies the chief effect of shuffling, the genetic diversification of gametes/offspring [1]. However, certain population genetic properties exhibit non-linear dependence on the rates of shuffling between loci. For example, theoretical studies have determined that certain allelic associations across loci can jointly increase in frequency only if their constituent loci shuffle their alleles at a rate below some critical threshold value. This has been shown for co-adapted gene complexes exhibiting positive fitness epistasis [139–141], ‘poison-antidote’ gamete-killing haplotypes involved in meiotic drive [10, 142–144], and associations between sex-determining genes and genes with sexually antagonistic fitness effects [52, 145–147]. Similarly, Hill-Robertson interference is effective only among loci that are tightly linked, shuffling their alleles at a rate below a small threshold (which depends on the effective population size and other parameters) [148]. In quantifying these various properties at an aggregate level, a more suitable measure would be the proportion of locus pairs that shuffle their alleles at a rate below the critical threshold. Such measures would be informative, for example, of which chromosomes are most likely to be co-opted as new sex chromosomes, or of how susceptible a species is to the invasion of ‘poison-antidote’ meiotic drive complexes (or, within species, which chromosomes are most likely to harbor such complexes), or of the average fitness reduction caused by Hill-Robertson interference. Calculating these threshold-based aggregate measures would require estimates of the average rates of shuffling for specific locus pairs. High-density linkage maps would therefore be especially promising for such measurements.

### Taking gene density into account

The calculation of *r̅*, as we have defined it, treats all regions of equal genomic length as of the same importance. However, the quantity we seek to measure is really the amount of shuffling of functional genetic elements—it is largely irrelevant if two functionless pseudogenes shuffle, for example. Therefore, correcting the calculation of *r̅* to take gene density into account is an important extension, a point already noted by Burt, Bell, and Harvey [26]. This is especially so if the distribution of genes is non-random along or across chromosomes. Accounting for the gene densities of different genomic regions would be easiest with sequence data, especially in species with well-annotated genomes. For species without well-annotated genomes, proxies of the gene density of genomic regions, such as euchromatin content or GC-richness, could be used. Accounting for gene density with cytological data would be more complicated, since it requires knowledge of the identity and orientation of the various chromosomes. At a crude resolution, this can be achieved with chromosome-specific FISH probes (and, if required, centromere localization). Finer-scale identification of different genomic regions in cytological spreads could be achieved using high-resolution FISH [149–151]. In any case, the usual problems would arise in deciding which parts of the genome are ‘functional’ (see [152, 153] for reviews of recent debates on this topic).

### Taking gene conversion into account

Another source of shuffling, in addition to crossing over and independent assortment, is gene conversion (GC) [154, 155]: the unidirectional copying of a DNA sequence tract from one chromatid (the donor) to a homologous chromatid (the acceptor), either in mitosis or meiosis. In SI Appendix, we show how to take both mitotic and meiotic GC into account in measuring *r̅*. In meiosis, chromosome pairing involves many points of interaction between homologous chromatids (~ 200 across all chromosomes in mouse spermatocytes [156]), each induced by a double-strand break on one of the chromatids. In most organisms, only a minority of these interactions result in crossovers [103, 157, 158], the majority instead resulting in non-crossovers. Both outcomes are associated with short tracts of GC, (e.g., ~ 100bp for non-crossover GCs and ~ 500bp for crossover GCs in mice [97, 156]). These tracts, though numerous, are too short to seriously affect the total amount of shuffling [for example, the contribution from meiotic GCs in mammals will be of order 10,000 times smaller than the contribution from crossovers (SI Appendix); in yeast, this figure is smaller—about 200 (SI Appendix)—owing to their longer GC tracts and smaller genomes], though they may have important local effects [154, 159]. Mitotic GC tracts result from homologous chromosome repair, are typically much longer than meiotic GC tracts (often of order 10kb; e.g., [160, 161]), and, if they occur in germline cell divisions leading to meiocytes, can lead to shuffling of maternal and paternal DNA in gametes (which should especially affect males, who have more germline cell divisions). Too little is presently known about the global frequency of these events for us to give a reasonable quantitative estimate of how much shuffling they cause.

## Acknowledgments

C.V. is grateful to Bob Trivers for impressing upon him the importance of crossover position for the amount of recombination. We are grateful to Anthony Geneva, David Haig, Arbel Harpak, Dan Hartl, Alison Hill, Thomas Lenormand, Pavitra Muralidhar, Naomi Pierce, and Martin White for helpful comments, and to Colin Donihue for help with figure preparation. Research of N.K. was supported by NIH GM-044794.

## Footnotes

1 Gene conversion also causes shuffling, but makes a negligible contribution, as discussed below.

2 Shuffling caused either by crossing over or independent assortment of homologs is referred to as ‘recombination’ in the population genetics literature, but ‘recombination’ has a different meaning—the breakage and rejoining of DNA molecules—in molecular biology. To avoid confusion, we use the unambiguous term ‘genetic shuffling’ (or just ‘shuffling’) throughout.

3 Given a gamete sequence and knowledge of the paternal and maternal origins of all chromosome segments, the realized value of r is precisely known—i.e., it is a parameter. If the maternal and paternal origins are not known, then *r̅* is a random variable, its realized value being different for different assortments of the possible parental origins of the various segments. We therefore calculate its expectation with respect to the distribution of these possible assortments. Similarly, when we observe crossovers along bivalent chromosomes at meiosis I, *r̅* in a resulting gamete is again a random variable, so we calculate its expected value, but this time with respect to the distribution of possible patterns of chromatid involvement in the resolution of crossovers at meiosis I, and the segregation pattern of chromatids thereafter.

4 In some species (e.g., tomato [114]), it has been reported that loops are longer in heterochromatin than in euchro-matin. While the data in Fig. S2 suggest that this is not generally the case, in such species appropriate adjustments would be needed to convert physical lengths to genomic lengths.

